# DNA lesions proximity modulates damage tolerance pathways in *Escherichia coli*

**DOI:** 10.1101/210237

**Authors:** Élodie Chrabaszcz, Luisa Laureti, Vincent Pagès

**Affiliations:** Team DNA Damage and Genome Instability, Cancer Research Center of Marseille, CRCM, Aix Marseille univ, CNRS, inserm, institut Paoli-Calmettes, 13009 Marseille, France

**Author notes:** These authors contributed equally to the paper as first authors (Élodie Chrabaszcz, Luisa Laureti).

## Abstract

The genome of all organisms is constantly threatened by numerous agents that cause DNA damages. When the replication fork encounters an unrepaired DNA lesion, two DNA damage tolerance pathways are possible: error-prone translesion synthesis (TLS) that requires specialized DNA polymerases, and error-free Damage Avoidance (DA) that relies on homologous recombination. The balance between these two mechanisms is essential since it defines the level of mutagenesis during lesion bypass, allowing genetic variability and adaptation to the environment, but also introducing the risk of generating genome instability. Here we report that the mere proximity of replication-blocking lesions that arise in *Escherichia coli*’s genome during a genotoxic stress, leads to a strong increase in the use of the error-prone TLS. We show that this increase is caused by the local inhibition of homologous recombination due to the overlapping of single-stranded DNA regions generated downstream the lesions. This increase in TLS is independent of SOS activation, but its mutagenic effect is additive with the one of SOS. Hence, the combination of SOS induction and lesions proximity leads to a strong increase in TLS that becomes the main lesion tolerance pathway used by the cell during a genotoxic stress.

## INTRODUCTION

The genome of all living cells is constantly exposed to DNA damaging agents that alter the chemical integrity of the DNA molecule. Cells possess efficient repair mechanisms such as Nucleotide Excision Repair (NER) or Base Excision Repair (BER) that are able to excise these DNA lesions, allowing replication to take place unhindered. However, some lesions might escape these repair mechanisms and block the progression of the replicative DNA polymerase during chromosomal replication. This leads to the generation of single stranded DNA (ssDNA) gaps downstream of the lesion: 1) when a lesion is located on the lagging strand, the lagging polymerase can be recycled or replaced at the next Okazaki fragment, leaving a ssDNA gap downstream of the lesion, without affecting the replication fork progression significantly; 2) when DNA damage occurs on the leading strand it is more problematic. Uncoupling of the leading - and lagging-strand synthesis have been reported both *in vitro* (1) and *in vivo* (2), allowing the lagging strand-synthesis to keep progressing for a certain distance. This implies that the replicative helicase (DnaB) keeps traveling for a distance, generating ssDNA downstream the lesion on the leading strand. The fate of the whole replication fork is still the subject of debate: one model sustains that DNA polymerization drastically slows down upon DNA damage, suggesting that the whole replisome stalls (3). On the other hand, earlier work has suggested that replication forks are able to skip over the lesion, leaving a gap that will be repaired later on (4). This model of “skipping” over the lesion has been recently corroborated by *in vitro* works showing that repriming can take place on the leading strand (5, 6), (7), and *in vivo* work showing that DNA Pol IV acts mostly at gaps left behind the replication fork, rather than at stalled replication forks (8). In both models, whether the replication fork stalls at the damage, or skips over the damage, ssDNA gaps are generated downstream the lesion both in the leading and lagging strand, and need to be filled (repaired) in order to achieve chromosomal replication. For this purpose, cells have evolved DNA Damage Tolerance (DDT) pathways: i) Translesion Synthesis (TLS) by which specialized DNA polymerases insert a nucleotide directly opposite the lesion with the risk of introducing a mutation (9); ii) Homology Directed Gap Repair (HDGR) (10) by which the single-stranded DNA gap generated downstream the lesion is filled by homologous recombination (11, 12). Additionally, we recently showed that cells were also able to tolerate the lesion by surviving on a single chromatid at the expense of losing the damaged chromatid (10) (Supplementary Figure 1). Our team has developed a system that allows to monitor both TLS and HDGR at the level of a single lesion inserted site-specifically in the chromosome of a living bacteria (13). Using this assay, we have previously shown that upon encounter with a single blocking lesion, TLS represents a minor pathway, while Damage Avoidance events (including HDGR and damaged chromatid loss) accounted for most of the survival (10, 13). We also showed that this partition between DDT pathways can be modulated by genetic factors. For instance, during a genotoxic stress, the induction of the SOS system leads to an increase in the expression level of specialized DNA polymerases favoring TLS over HDGR (14). Also, when the homologous recombination machinery is impaired, the decrease in HDGR is accompanied by an increase in TLS (15). Besides the modulation of the actors of both pathways, either natural (SOS induction) or artificial (genes deletions or mutations), could a naturally occurring perturbation in the structure of the replication fork affect the partition in lesion tolerance? In this work we raised the question of what happens to the replication fork structure when lesions are present in opposite strands (an event occurring frequently during a genotoxic stress), and showed that this event affects lesion tolerance by preventing Homologous Recombination (HR) and favoring TLS..

## RESULTS

### Lesion proximity increases TLS

In order to study the impact of lesions proximity on DDT pathways, we constructed integrating vectors that harbor one AAF (N2-AcetylAminoFluorene) lesion in each strand. These vectors were introduced at a specific locus in the chromosome of a living bacteria using a recently developed system that allows to monitor both TLS and HDGR at the damaged sites (13, 19) (Supplementary Figure 1). We initially chose a distance of 1.8kb to separate the two lesions in order to mimic a realistic genotoxic stress. Indeed, such density of one lesion every 1.8kb corresponds to ~2500 lesions per chromosome which is equivalent to UV irradiation of ~50J.m-2 (4, 21). In our original vector design, the AAF lesion was located in the lacZ reporter gene (Figure 1A). We kept this lesion in the same locus and added the second lesion in the chloramphenicol resistance gene (cat) on the opposite strand (Figure 1B). Both AAF lesions are located in the NarI mutation hotspot where Pol V mediates error-free TLS (TLS0) and Pol II mediates-2 frameshift TLS (TLS-2) (22). A 3-nucleotide loop opposite to each lesion allows to monitor HDGR events.

**Figure 1:**
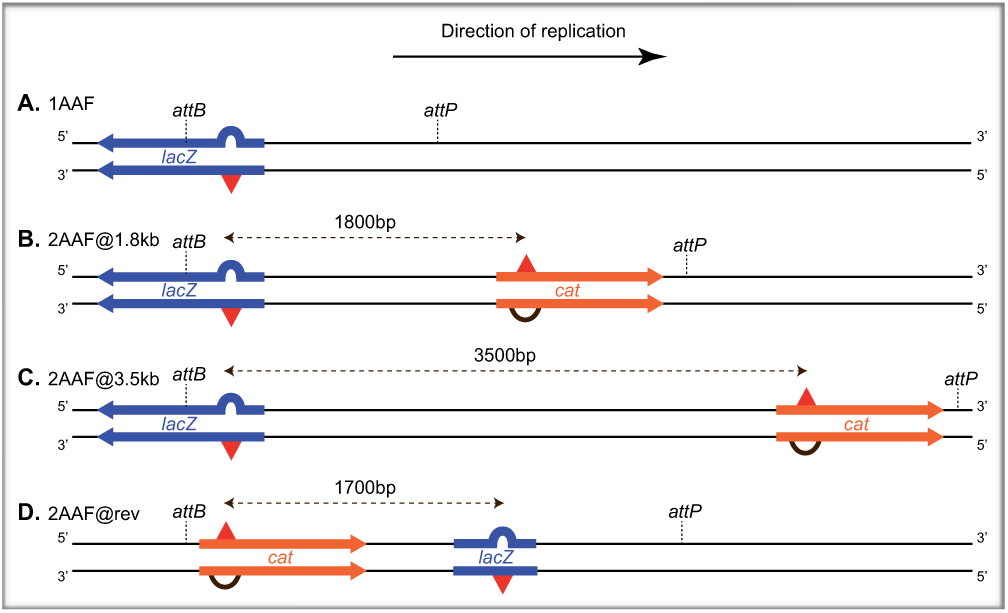
Map of the lesions once integrated in the bacterial genome between phage lambda attB and attP recombination sites. A: a single AAF lesion in the *lacZ* gene. B and C: 2 lesions are separated by 1.8 and 3.5kb. The first lesion encountered by the replication fork is located on the leading strand (*lacZ* gene) while the second lesion encountered is on the lagging strand (*cat* gene). D: reverse configuration where the lesions are separated by 1.7kb, the first lesion encountered by the replication fork is located on the lagging strand while the second lesion is on the leading strand.

When monitoring TLS in our parental strain (uvrA, mutS deficient to prevent repair of the lesion and of the strand markers) (Figure 2) we observe that the presence of a second lesion at a distance of 1.8kb strongly increases TLS at the original (lacZ) lesion site. We indeed observe a 3-fold increase in Pol V TLS, and a 2.5-fold increase in Pol II TLS (Figure 2). Concomitant with this increase in TLS, we observe a strong decrease in survival (Figure 3A). A single lesion doesn’t induce any toxicity and is mostly tolerated by HDGR, while damaged chromatid loss accounts for a small fraction of survivors and TLS is a minor event representing ~2% of the surviving cells. The presence of the second lesion appears to be highly toxic for the cell since the viability drops to ~25%. The massive loss of viability is partly due to the fact that damaged chromatid loss is no longer possible since both chromatids are damaged, but mostly to a strong decrease in HDGR. We hypothesized that this inhibition of HDGR was caused by the overlapping of single stranded DNA gaps generated at the opposite lesions (Figure 5B). This suggests that the replicative helicase had opened the DNA duplex over a length of 1.8kb and that if repriming had occurred, it would have had occurred beyond 1.8kb on the leading strand, and that the Okazaki fragment preceding the lesion had initiated before 1.8kb on the lagging strand (Figure 5B). In this situation, no dsDNA substrate is available opposite any of the lesions to allow RecA nucleofilament invasion (D-loop) that is the initial step of HDGR (23, 24). This strong decrease in HDGR is leading to the observed increase in TLS (Figure 2). This is consistent with our previous observation that TLS increased when HR was impaired genetically by mutations in recA or inactivation of recF or recO (15). Because of the combined increase in TLS and decrease in HDGR, total TLS events (including TLS at both AAF sites) represent now more than 40% of the surviving cells (as compared to 2% for a single lesion).

**Figure 2:**
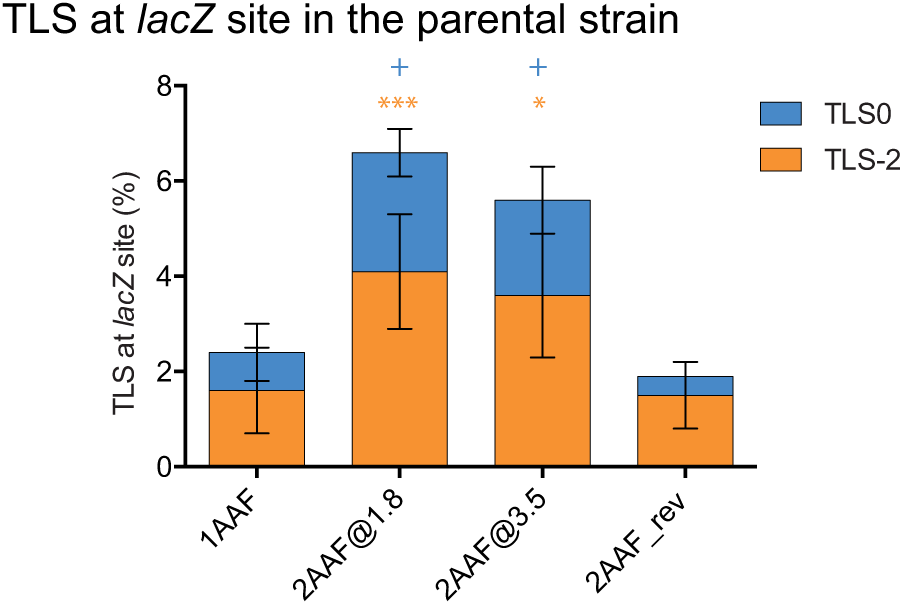
TLS events at the original *lacZ* locus in the parental strain (*uvrA mutS*) for the different lesion configurations. Pol V TLS is error-free (TLS0), while Pol II TLS leads to a −2 frameshift (TLS-2). The data represent the average and standard deviation of at least 3 independent experiments. Unpaired t-test was performed to compare TLS values from the integration of 2 lesions to the integration of a single lesion. For TLS0: +P<0.05. For TLS-2: *P<0.05; ***P>0.0005

**Figure 3:**
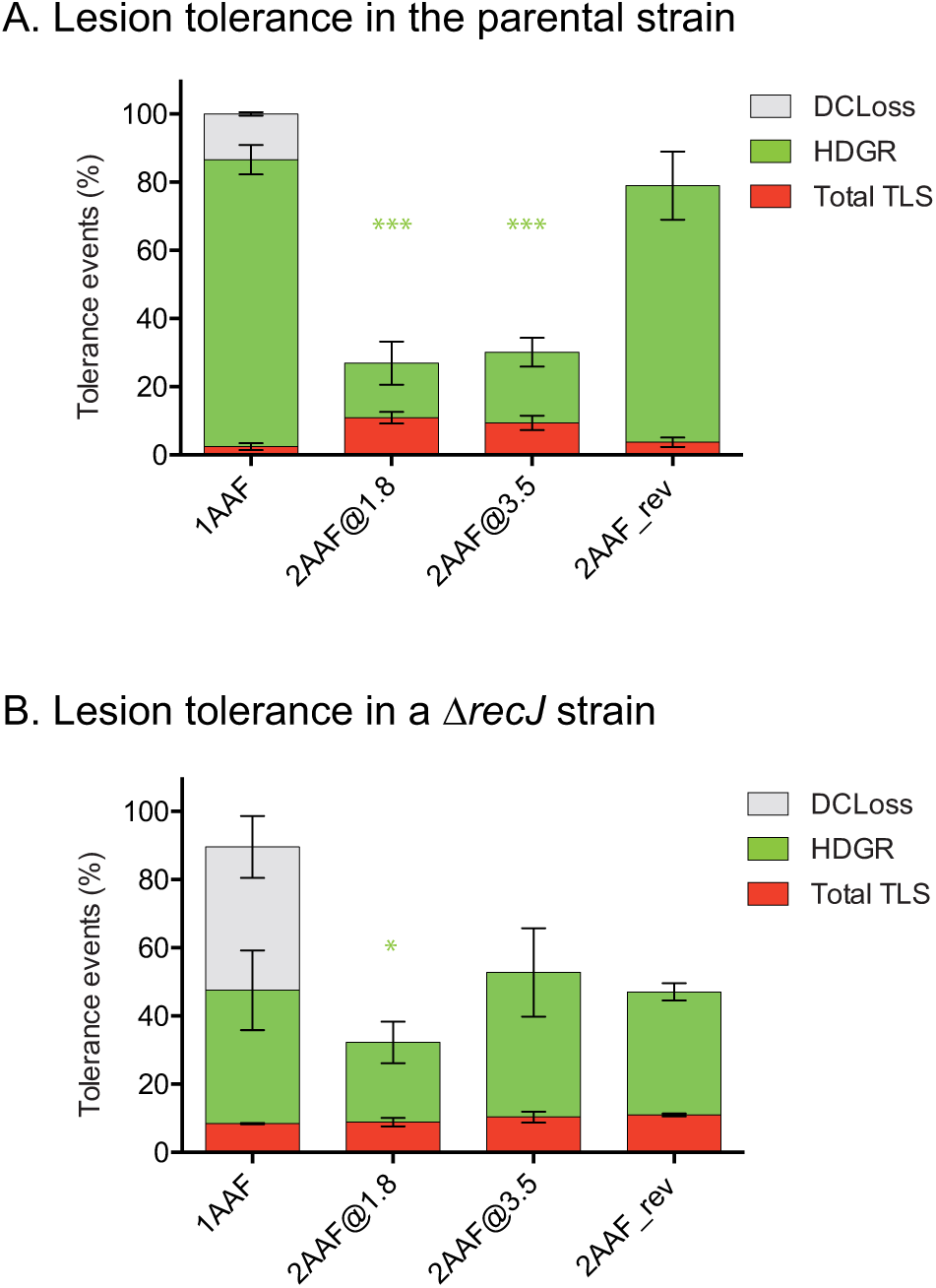
Partitioning of the events allowing the cell to survive the damages in the parental strain (A) and in the ∆recJ strain (B). Tolerance events (Y axis) represent the percentage of cells able to survive in presence of the integrated lesion(s) compared to the lesion-free control. Total TLS (or HDGR) represents the percentage of cells that achieved TLS (or HDGR) at the lacZ site and/or at the cat site when present. Damaged chromatid loss (DC Loss) is only observed for the single lesion construct. The data represent the average and standard deviation of at least 3 independent experiments. Unpaired t-test was performed to compare HDGR values from the integration of 2 lesions to the integration of a single lesion. *P<0.05; **P<0.005; ***P>0.0005

### 5' end resection by RecJ exonuclease modulates the length of ssDNA gaps

We then reasoned that by increasing the distance between the two lesions, we should avoid this overlap of single strand gaps and restore some viability. We constructed and integrated a new vector where both AAF lesions are now 3.5kb apart (Figure 1C).

Compared to the previous situation where lesions were 1.8kb apart, neither TLS (Figure 2) nor HDGR or survival (Figure 3A) were significantly different. It appears therefore that even at 3.5kb, the gaps generated at the lesion are still overlapping, inhibiting HDGR and favoring TLS. It is unlikely that repriming events on the leading strand would occur beyond 3.5kb, since the gaps observed after UV irradiation have been estimated to be in the range of ~1-2kb (25). Similarly, on the lagging strand, Okazaki fragments are expected to be shorter than 3kb (26). Therefore, both in the leading and lagging strand, one can expect dsDNA on the strand opposing the lesion, and HR should be observed. Since we didn’t observe any recovery in HDGR when lesions are 3.5kb apart, we hypothesize that the 5' end of the reprimed fragment or of the Okazaki fragment preceding the lesion could be resected by a 5'>3' exonuclease, creating overlapping ssDNA regions that would prevent HDGR.

Daughter strand gap repair occurs through the RecF pathway where RecFOR mediates the loading of RecA protein onto SSB-coated ssDNA (27). Belonging to the RecF pathway, it was proposed that RecQ helicase and RecJ nuclease participate to the process by widening the gaps (28-30). RecJ exonuclease possesses a 5'3' polarity (31) that combined with the action of RecQ helicase can resect the newly synthesized DNA at the previous Okazaki fragment on the lagging, or at the reprimed fragment downstream the lesion on the leading strand if repriming has occurred. Consistent with this role of RecJ, following the introduction of a single G-AAF lesion, the inactivation of recJ (comparison of 1AAF between Figure 3A and Figure 3B, and Figure 4) leads to a strong decrease in HDGR that is mostly compensated by damaged chromatid loss events, and also accompanied by an increase in TLS as previously observed for recF or recO mutants (15). When a second lesion is present at 1.8kb, recJ inactivation has no effect on the repartition of DDT events. However, when a second lesion is present at 3.5kb, recJ inactivation restores some viability by increasing HDGR (compare 2AAF@3.5 between Figure 3A and 3B) to the level observed with a single lesion. This result supports our model where HDGR is prevented by the overlapping of opposite single strand gaps: in the presence of RecJ exonuclease, gaps are extended beyond 3.5kb causing the ssDNA-gaps to overlap and preventing HDGR to occur; in the absence of RecJ, repriming events that occur before 3.5kb are not resected and allow HDGR (Figure 5C). This observation with only 2 DNA lesions artificially introduced on the genome would imply that RecJ has opposite effects depending on the density of lesions. At low lesion density, DNA resection by RecJ would expand the gaps and increase the efficiency of HDGR, whereas at high lesion density, the gaps expansion would lead to their overlapping that would prevent HDGR. That is indeed what has been observed by Courcelle et al. who compared UV-survival of a recJ to a WT strain (32): at low doses, the recJ strain is more sensitive than the WT strain showing the requirement of RecJ for HDGR. At higher doses, the recJ strain becomes more resistant than the WT strain showing the deleterious effect of RecJ when lesion density is higher. Interestingly, the UV dose where the effect of RecJ switches is in the range of 60J.m-2, which corresponds to a lesion density that is in the same range as our experimental setup.

The decrease in HDGR observed in the recJ strain in the presence of a single AAF lesion supports the model by which RecJ widens the ssDNA gaps generated downstream of a DNA lesion. Interestingly, while RecJ has been mostly described as acting at the 5' terminus of an Okazaki fragment on the lagging strand (33), we observe the same effect whether the lesion is located on the lagging or on the leading strand (Figure 4), indicating that RecJ can act with similar efficiency both at a repriming event that occurred on the leading strand, and at an Okazaki fragment on the lagging strand. This observation favors the model where repriming occurs on the leading strand as suggested by in vitro data (5, 6).

**Figure 4:**
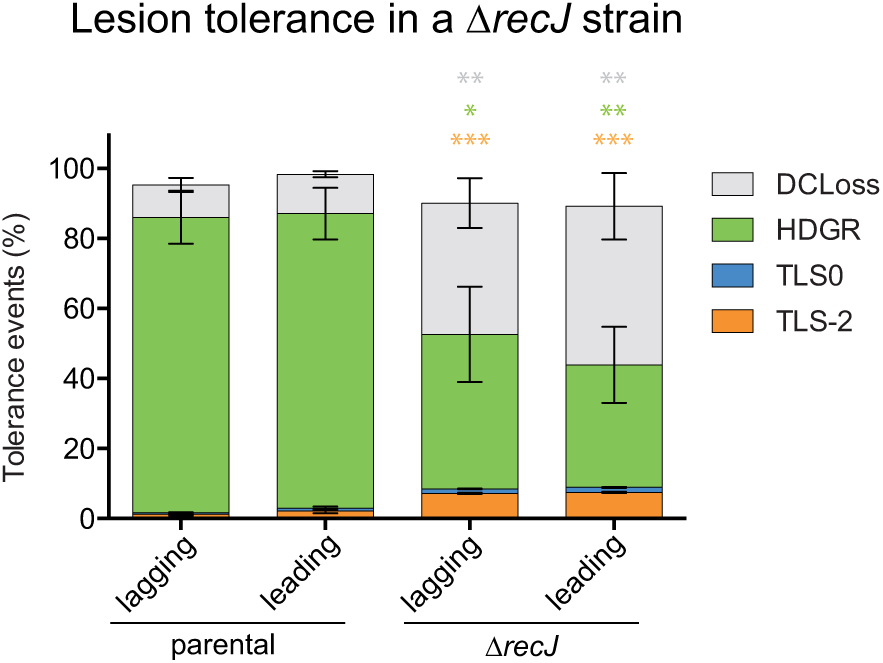
Partitioning between TLS, HDGR and Damaged chromatid loss (DC Loss) in the presence of one single AAF lesion inserted in the parental strain and in the *recJ* deficient strains. The lesion has been inserted in either the leading or the lagging strand of *E. coli* chromosome. Tolerance events (Y axis) represent the percentage of cells able to survive in presence of the integrated lesion compared to the lesion-free control. The data represent the average and standard deviation of at least three independent experiments. No significant difference is observed in the level of HDGR whether the lesion is inserted on the lagging or leading strand. The data represent the average and standard deviation of at least 3 independent experiments. Unpaired t-test was performed to compare values from parental strain to the ∆*recJ* strain. *P<0.05; **P<0.005; ***P<0.0005

### The initial gap generated downstream the lesion is in the range of 1.8 to 3.5kb

It is interesting also to note that in the recJ strain (Figure 3B), the survival is much lower with 2 lesions since damaged chromatid loss that accounted for ~50% of survival with a single lesion is no longer possible (both chromatids being damaged). However, the level of HDGR is similar when 2 lesions are 3.5kb apart and when one lesion is isolated. This suggests that in the absence of resection (∆recJ), the amount of ssDNA downstream of the lesions is similar in both situations, and therefore that the gap formed downstream a single lesion is shorter than 3.5kb (before resection). Together with the previous observation that HDGR is strongly inhibited when the lesions are 1.8kb apart (whether RecJ is present or not), we can conclude that the gap generated downstream a lesion is in the range of 1.8 to 3.5kb. This size of gap (in the absence of expansion by RecJ) seems insufficient to promote efficient homologous recombination since recJ deletion leads to a strong decrease in HDGR at a single lesion. Further studies will be required in order to determine how far the gap is extended by RecJ in order to achieve WT-levels of HDGR.

### Single-strand gaps overlapping prevents HDGR

To further challenge our model, we constructed a vector with two lesions in a reverse configuration where the first lesion encountered by the replication fork is located in the lagging strand, and the second lesion, positioned 1.7kb downstream is located in the leading strand (Figure 1D). In this configuration, no matter how far the fork unwinds or where the repriming occurs, or whether gap widening occurs, the ssDNA gaps generated downstream of each lesion are not overlapping and therefore, one does not expect HDGR inhibition (Figure 5D). Integration of this vector indeed shows a high level of HDGR similar to the one observed with a single lesion, despite the proximity of the two lesions (Figure 3A). In addition, the TLS level at the initial lacZ locus is not increased by the proximity of the second lesion (Figure 2). This observation rules out the possibility that the increase in TLS observed with the constructions 2AAF@1.8 and 2AAF@3.5 was due to further SOS induction by the additional lesion, and confirms our hypothesis that TLS increase is the result of HDGR inhibition by overlapping ssDNA gaps.

**Figure 5:**
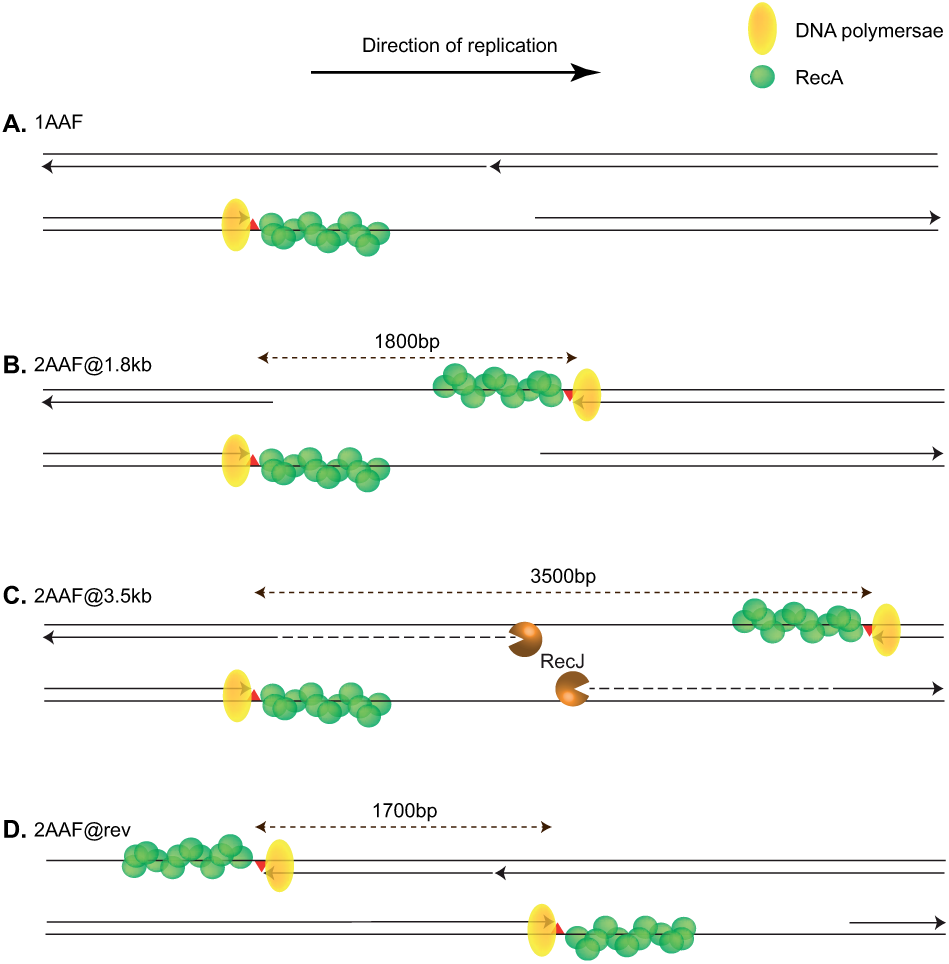
Structural configuration during lesion bypass. A: when one single lesion is encountered in the leading strand, a ssDNA gap is generated downstream the lesion. The gap can be filled by HDGR or TLS. B: when 2 lesions distant from 1.8kb are present on opposite strands, the two ssDNA regions generated on opposite strands prevent HDGR and favor TLS. C: when the 2 lesions are separated by 3.5kb, 5'-end resection by RecJ leads to overlapping ssDNA gaps that prevent HDGR and favor TLS. In the absence of RecJ, ssDNA gaps are no longer overlapping and HDGR is possible. D: in the reverse configuration, when the first lesion encountered by the replication fork is located on the lagging strand, no overlapping single strand gaps occurs and HDGR level is similar to the one of a single lesion. In all situations, repriming is possible on the leading strand and was represented. But whether repriming occurs in vivo and where it occurs is still subject of debate (see text).

### Additional effects of lesions proximity and homologous recombination inhibition

In our model, the mere proximity of DNA lesions leads to a structural inhibition of homologous recombination that in turn favors Translesion synthesis. This structural inhibition is distinct from the previously observed genetic inhibition of HDGR that was caused by the inactivation of recF or recO (15). RecFOR is known to mediate the loading of RecA onto SSB-coated ssDNA. Inactivation of recF delays the loading of RecA and therefore reduces the efficiency of gap repair: when introducing a single AAF lesion in a ∆recF strain, we observe a decrease in HDGR accompanied by an increase in damaged chromatid loss and a 4-fold increase in TLS-2 (Figure 6A) compared to the parental strain (Figure 2 and 3A). The addition of a second AAF lesion in the ∆recF strain further decreases HDGR, and leads to an additional increase of TLS, showing that the two mechanisms are indeed distinct: recF inactivation reduces RecA loading efficiency, whereas lesion proximity sequesters the substrate for HR (dsDNA). Both effects are additive and lead to an overall >7 fold increase in TLS-2 (Supplementary Figure 3 clearly shows the additional effect of recF deletion and lesion proximity on TLS).

### Additional effects of lesions proximity and SOS induction

We also show that the increase in TLS is independent of SOS activation or of any modulation of genetic factors. Such a constraint due to the proximity of DNA lesions may occur naturally and quite frequently during a genotoxic stress, and allows cells to modulate their DNA damage response by favoring TLS. Strong genotoxic stresses are also known for inducing the SOS response that favors TLS by increasing the expression of specialized polymerases. By introducing one or 2 lesions in a lexA deficient strain where the SOS system is constitutively induced, we show that the two mechanisms are indeed additive (Figure 6B): the SOS induction leads to a ~5-fold increase in the use of the TLS pathway, and the proximity of the DNA lesions leads to an additional ~2-fold increase in the use of TLS. Overall, error-prone TLS accounts for ~90% of survival when lesions are close by and SOS is induced.

**Figure 6:**
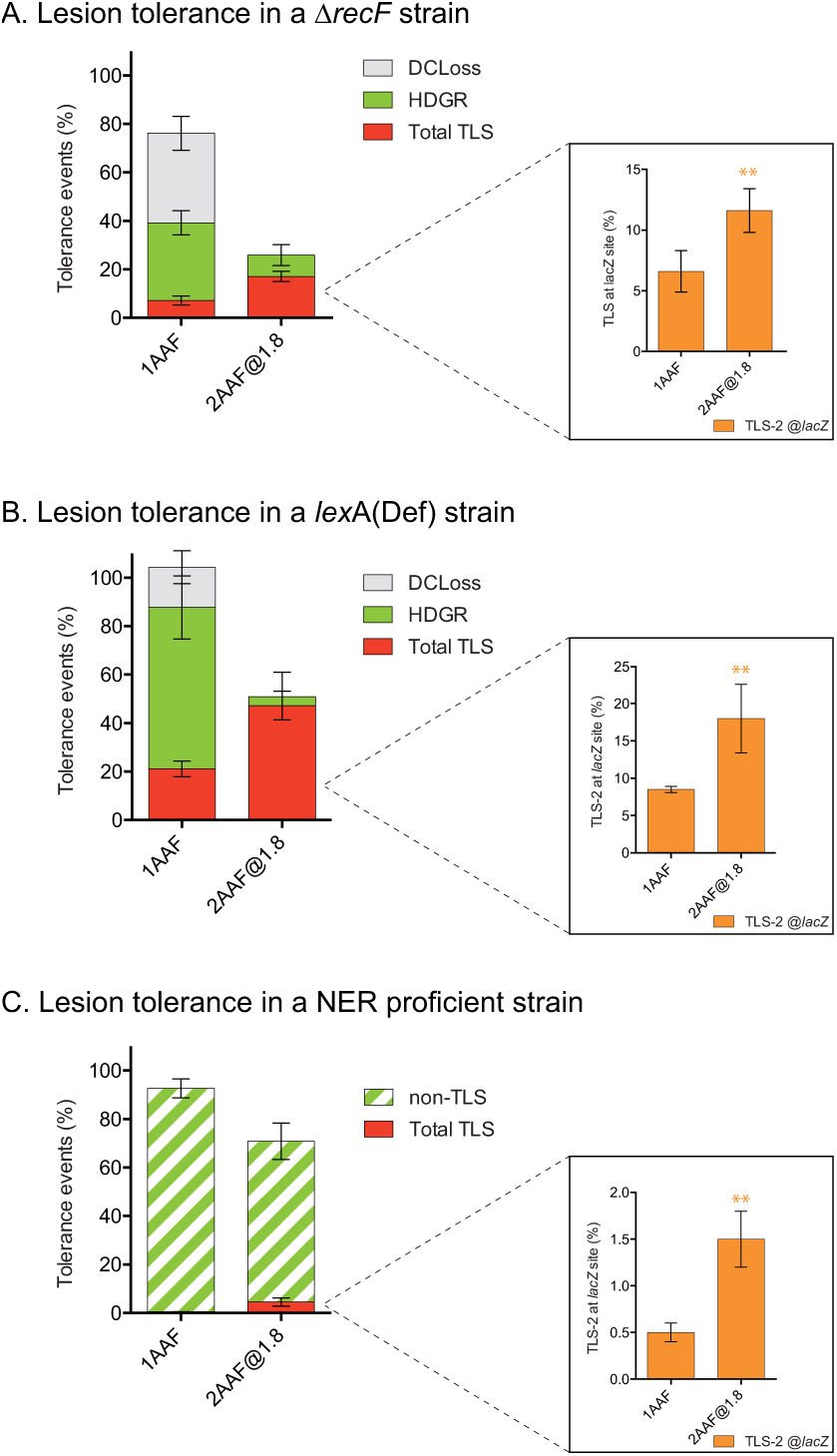
Partitioning of DDT pathways at one single AAF lesion and at 2 AAF lesions on opposite strands at 1.8kb distance A: in a recF strain. B: in a lexA(Def) strain where the SOS system is constitutively induced. C: in a strain proficient for Nucleotide Excision Repair (uvrA+). The inserts represent Pol II mediated TLS-2 at the lacZ site only, to allow direct comparison of mutagenic TLS between one and two lesions. The data represent the average and standard deviation of at least 3 independent experiments. Unpaired t-test was performed to compare TLS values from the integration of 2 lesions to the integration of a single lesion. *P<0.05; **P<0.005; ***P<0.0005

### Lesions proximity favors TLS in NER proficient cells

Finally, in order to extrapolate this observation to a wild-type context where lesions can be repaired, we introduced the lesions in the genome of a Nucleotide Excision Repair proficient (NER+) strain (Figure 6C). The level of TLS at a single AAF is much lower in the NER+ strain compared to the NER- strain (Figure 2) due to the fact that the lesion is frequently repaired before TLS could occur. When integrating two lesions, we observe a strong recovery in the survival in the NER+ compared to the NER- strain, that is mostly due to repair (that cannot be distinguish from HDGR in our system). Despite this high level of repair, the structural effect due to the lesions proximity at unrepaired lesions persists and leads to a strong increase in TLS compared to the single lesion.

## DISCUSSION

In this work, we show that the mere proximity of DNA lesions on opposite strands leads to an inhibition of homologous recombination, and to a concomitant increase in TLS. The inhibition of homologous recombination is due to the overlapping of ssDNA regions that are generated at each lesion. In the lagging strand, the replicative DNA polymerase blocked at the lesion is recycled at the next Okazaki fragment, generating a ssDNA region downstream of the lesion. In the leading strand, the helicase progression uncovers a ssDNA region that we estimate in the range of 1.8 to 3.5kb. Whereas the replication fork stalls there or a repriming event allows it to keep going forward is still subject of debate. Such repriming events have been showed in vitro (5, 34), and recent in vivo data also suggest that gaps are left behind the fork (8). Our present data showing that gap widening by RecJ (a 5'3' exonuclease) favors HDGR at an isolated lesion whether the lesion is located on the leading of the lagging strand (Figure 4) are also in favor of this model.

We also show that even when lesions are rather far apart (3.5kb), the increase in TLS is still observed due to the expansion of ssDNA gaps by RecJ.

The inhibition of homologous recombination due to the overlap of ssDNA is distinct from the genetic inactivation of recF, and the effect on TLS increase is additive with the one of recF inactivation. Similarly, the effect on TLS increase is independent of SOS induction, but again additive with it. Finally, we show that the effect of lesions proximity (initially revealed in a NER deficient background) persists in a NER+ strain despite the fact the majority of DNA lesions are repaired in this background.

Overall, this phenomenon appears like a structural mechanism that acts independently of genetic factors and other mechanisms that are known to regulate the balance between error-free and error-prone lesion tolerance mechanisms. It allows the cell to respond to a genotoxic stress by enhancing the error-prone translesion synthesis: the accumulation of mutations resulting form TLS will generate the genetic diversity required for the cell to adapt and survive such conditions. Since the basic mechanism of homologous recombination is conserved among species, this inhibition by lesions proximity and the concomitant increase in TLS might be a new paradigm for genotoxic stress induced mutagenesis that could potentially be applied to all organisms.

## MATERIAL AND METHODS

### Plasmid construction

Vector harboring a single lesion or dual lesions are constructed using the gap-duplex method as previously described. Supplementary table 1 shows all the plasmids used in this study.

pEC29 and pEC30 are derived from previously described plasmids pVP143 and pVP144 (6). The chloramphenicol resistance gene (*cat*) and its promoter have been added in the opposite orientation with respect to the lacZ gene, in order to serve as a reporter gene where is introduced the second lesion. The 5' end of the cat gene has been modified by site directed mutagenesis in order to allow insertion of the AAF-modified oligonucleotide (plasmid pEC30) and the Nar+3 strand marker on the opposite strand (pEC29).

pEC37 and pEC38 are modified versions of pEC29 and pEC30 where a 1.7kb spacer (mCherry and GFP genes without their promoter) have been inserted between the two reporter genes (*lacZ* and *cat*) in order to increase the distance between the two lesions.

pEC45 and pEC46 are modified versions of pEC29 and pEC30 where the cat gene was cloned on the other side of *lacZ* gene regarding the attL integration site. A transcription termination site was added between *lacZ* and cat genes to avoid any potential interference between transcription and DDT events at the cat lesion. The integration of the pEC45/ pEC46 duplex doesn’t reconstitute a functional *lacZ* gene (see Figure 1D), so TLS events are monitored by sequencing.

All six vectors contain the following characteristics: the R6K replication origin that allows the plasmid replication only if the recipient stain contains the pir gene, the ampicillin resistance gene that allows selection of integrated colonies, the chloramphenicol resistance gene as reporter for TLS at one lesion, and the 5' end of the lacZ gene in fusion with the attL recombination site of phage lambda. The P’3 site of attL has been mutated (AATCATTAT to AATTATTAT) to avoid the excision of the plasmid once integrated. These vectors are produced in strain EC100D *pir-116* (from Epicentre Biotechnologies - cat# EC6P0950H) in which the *pir-116* allele supports higher copy number of R6K origin plasmids.

### Strains

All strains are derived from FBG151 and FBG152 (9). After integration of the vector in the attR site, the lesion at the *lacZ* site is located on the lagging strand in FBG151 and its derived strains, and on the leading strand in FBG152 and its derived strains. Gene disruptions were achieved by the one-step PCR method (10). The following FBG151 and FBG152 derived strains were constructed by P1 transduction (Supplementary table 2).

### Monitoring DDT events

Competent cells preparation and integration of lesion-containing vectors were conducted as previously described (6, 11). Briefly, a non-damaged control vector and the lesion(s) containing vector are transformed together with an internal standard plasmid (pVP146) in electrocompetent cells expressing the *int-xis* gene of phage lambda. After a 45 minutes incubation period, cells are plated on LB agar media containing antibiotic and X-gal indicator. Survival is calculated by the ratio of colonies resulting from the integration of the damaged vector over the non-damaged control, corrected by the transformation efficiency of the internal standard plasmid (pVP146).

For the single lesion containing vectors, Pol V mediated TLS0 and Pol II mediated TLS-2 were measured by counting blue colonies after integration of pVP141/142 and pVP143/pVP144 duplexes respectively (6). HDGR and damaged chromatid loss was assessed by monitoring blue and white colonies after the integration of pLL1/pLL7 as previously described (2, 12).

For the dual lesions containing vectors pEC29/ pEC30 and pEC37/pEC38, TLS-2 at lacZ site was measured as blue colonies on X-gal indicator plate and TLS-2 at cat site was measured as chloramphenicol resistant colonies. All events (including TLS-2) for these plasmids and for pEC45/ pEC46 were monitored by Sanger sequencing after whole colony PCR amplification of the damaged region. Chromatogram analysis allows to visualize events that occurred on both strands of each damaged site (Supplementary Figure 1). Figure 2 represents all TLS events that occurred at the *lacZ* site regardless of what has happened at the other site. In Figures 3A-B and Figure 6, “total TLS” represents the percentage of cells that survived using TLS at lacZ and/or at Cm site. HDGR represents the percentage of cells that survived using HDGR at lacZ and/or at Cm but didn’t use TLS. Detail of the tolerance events at both sites are presented in supplementary table3.

Integrations were performed in FBG151 and FBG152 derived strains allowing lesions at the *lacZ* site and at the *cat* site to be alternatively on the lagging or leading strand. Since no difference was observed for the 2 orientations, the graphs and table represent the average of integration events obtained in both orientations.

## Acknowledgments

this work was supported by Agence Nationale de la recherche (ANR) Grant GenoBlock ANR-14-CE09-0010-01. We thank Mauro Modesti for critical reading of the manuscript. We thank Ryan Muldoon for proofreading the article.

